# Pollination services to crops of watermelon (*Citrullus lanatus*) and green tomato (*Physalis ixocarpa*) in the coastal region of Jalisco, Mexico

**DOI:** 10.1101/2024.03.18.585619

**Authors:** Oliverio Delgado-Carrillo, Silvana Martén-Rodríguez, Diana Ramírez-Mejía, Samuel Novais, Alexander Quevedo, Adrian Ghilardi, Roberto Sayago, Martha Lopezaraiza-Mikel, Erika Pérez-Trujillo, Mauricio Quesada

**Affiliations:** Laboratorio Nacional de Análisis y Síntesis Ecológica, Escuela Nacional de Estudios Superiores Unidad Morelia, Universidad Nacional Autónoma de México, Morelia, Michoacán, Mexico; Instituto de Ecología, Universidad Nacional Autónoma de México, Ciudad de México, México; Comisión Nacional para el Conocimiento y Uso de la Biodiversidad, Ciudad de México, México; Red de Interacciones Multitróficas, Instituto de Ecología A.C., Xalapa, Veracruz, México; Centro de Investigaciones en Geografía Ambiental, Universidad Nacional Autónoma de México, Morelia, Michoacán 58190, México; Facultad de Desarrollo Sustentable, Universidad Autónoma de Guerrero, Tecpán de Galeana, Guerrero, Mexico; Facultad de Biología, Universidad Michoacana de San Nicolas de Hidalgo, Morelia, Michoacán, Mexico; Instituto de Investigaciones en Ecosistemas y Sustentabilidad, Universidad Nacional Autónoma de México, Morelia, Michoacán, México

## Abstract

Bees play a pivotal role as pollinators in crops crucial for human consumption. However, the global decline in bee populations poses a significant threat to pollination services and food security worldwide. The loss and fragmentation of habitats due to land-use change are primary factors contributing to bee declines, particularly in tropical forests facing high deforestation rates. Here we evaluate the pollination services on crops of watermelon (*Citrullus lanatus*) and green tomato (*Physalis ixocarpa*) in the Tropical Dry Forest of Jalisco, Mexico during 2008, and 2014 to 2017. We describe pollinator community composition and visitation frequency and evaluate pollinator contribution to plant reproductive success and degree of pollinator dependence for each crop species. We also assess how landscape configuration influences the abundance and richness of pollinators, and we use the model Integrated Valuation of Ecosystem Services and Tradeoffs (InVEST) to map and value the pollination service in both crops. Our findings reveal that the main pollinator of both crops was *Apis mellifera*, one of the few abundant pollinators in the study region during the dry season, when watermelon and green tomato are cultivated. Results revealed that in the absence of pollinators, watermelon yielded no fruits, while green tomato experienced a 65% reduction in production. For green tomato, fruit set was positively correlated with pollinator abundance. A positive association between forest cover and total pollinator abundance was observed in green tomato in 2008, but not in watermelon. We also found a positive relationship between the abundance of bees predicted by the InVEST model and the abundance of bees observed in green tomato flowers in 2008. In the study region, green tomato and watermelon rely on pollinators for fruit production, with honeybees (from feral and managed colonies) acting as the most importante provider of pollination services for these crops. Consequently, the conservation of natural protected areas is crucial for sustaining pollinators and ensuring food security.

## Introduction

Approximately 70% of major food crops worldwide rely on animal pollinator services to ensure crop production (1). This value is even higher in tropical regions; for instance, in Mexico, approximately 85% of crop species depend on animals for pollination (2). Bees are the most important group of flower visitors in pollinator-dependent crops, accounting for 62% of the visits to crop flowers; the remaining 38% consists mainly of other insect groups such as flies, butterflies, moths, wasps and beetles (1,3,4). The commercial European bee, *Apis mellifera*, is the most widely used species to ensure the pollination of ∼90 commercial crop species worldwide (4–7); however, wild bees are also important contributors to crop pollination, representing 23% of floral visitors worldwide (4,8–11).

Native plant species share evolutionary histories with their native pollinators, so it is reasonable to expect that native pollinators must be more efficient at providing pollination services than introduced pollinators in native crops, as has been demonstrated in various native crop species (2,8,11). For example, native pollinator are the most effective pollinators of blueberries (*Vaccinium corymbosum*) and squash (*Cucurbita* spp) grown within their areas of origin (12–15). Nonetheless, exotic pollinators are as effective as native pollinators in other crop species, as is the case of managed and native *Bombus* species in avocado, sweet pepper and tomatoes crops (16–18). *Apis mellifera* and some exotic *Bombus* species have become important pollinator for certain crops, both native or non-native (17,19–22), and feral populations of *A. mellifera* have established in natural ecosystems across the Neotropics, being omnipresent in natural and agricultural ecosystems (21). The action of both native and exotic pollinators is necessary to achieve full seed set in some species (e.g., 23–25). Therefore, the persistence of native and non-native pollinators is essential to ensure pollination services for crops worldwide.

Despite the fundamental role that commercial and native bees play in crop pollination and global food security, several native species of bees are threatened and populations of *A. mellifera* are experiencing declines. At least, eight major drivers of global pollinator declines have been described, including changes in land cover and configuration, land management, pesticide use, climate change, pest and pathogens, pollinator management, invasive species, and genetically modified organisms (26,27). Agricultural environments represent low quality habitats for pollinators as these systems do not provide sufficient resources for bee survival, nor suitable biophysical conditions for their persistence and reproduction (28,29). Furthermore, agricultural intensification negatively affects the yield of pollinator dependent crops (30). However, natural or semi-natural habitats near agricultural fields represent important refuges for pollinators as they offer nesting and foraging sites (31–33). Moreover, the shape and size of natural habitat patches, and their proximity to crops can affect the abundance and richness of pollinators (29,34,35). For instance, proximity to natural habitats can increase pollinator visitation rates (36) and improve crop yields, seed set and progeny quality (28,34,36–38). Hence, natural habitats can be considered key reservoirs of floral visitors that may contribute significantly to the pollination services required by certain crop species (28,39–41).

Regardless of the enormous importance of natural forests to the persistence of pollinators, anthropogenic activities pose a continuous threat to natural ecosystems. Evidence indicates that tropical pollinators, particularly bees, are very susceptible to land-use change and agricultural intensification (29,42). However, the relevance of natural ecosystems for the maintenance of wild pollinators and the pollination services they provide is little known for many tropical agroecosystems. One of the most threatened tropical ecosystems is the Tropical Dry Forest (TDF), which covers more than 40% of the world’s tropical forests (43). Only a third of TDF occurs in well-preserved continuous forest fragments, and the rest is found in fragmented and degraded forests (44). In Mesoamerica, TDF is among the most disturbed and least conserved ecosystems (45–48). TDF used to cover nearly one third of the Mexican territory, but it has become a highly threatened ecosystem, underrepresented in protected areas, and poorly studied (49). Crops that are cultivated in this ecosystem include watermelon (*Citrullus lanatus*), green tomato (*Physalis ixocarpa*), chili (*Capsicum annuum*), cucumber (*Cucumis sativus*), chayote squash (*Sechium edule*), papaya (*Carica papaya*), mango (*Manguifera indica*), and tomato (*Solanum lycopersicum*); thus, studying the contribution of protected areas to pollination is crucial for the maintenance of crop pollinator services and the food security of many rural communities.

Here, we assessed pollination services for two economically important crops: watermelon (*Citrullus lanatus*) and green tomato (*Physalis ixocarpa*) in the coastal region of Jalisco, Mexico. The specific goals of the study were: (i) to describe the community composition of pollinators associated with both crops, (ii) to evaluate floral visitation patterns by different pollinators across the day; (iii) to determine the efficiency of floral visitors by evaluating visitation rate, pollen on insect bodies, and the contribution of honeybees and native bees to fruit and seed set (iv) to determine the level of pollinator dependence for the two crops, (v) to evaluate the influence of landscape configuration on pollinator abundance and richness of both crops; and (vi) to map pollination services provided by bees to these crops in the region. We hypothesized that well-preserved patches of tropical dry forest play a key role in providing nesting sites and floral resources to bee pollinators; therefore, we expected their abundance and richness across mosaics of pollinator-dependent crops and tropical dry forests. This in turn will enhance crop pollination services overall.

## Material and methods

### Study area

We conducted the study in the municipalities of La Huerta, Cihuatlan, and Tomatlán in the southwestern coast of Jalisco, Mexico (Fig 1). Natural vegetation in this region is dominated by Tropical Dry Forest (∼56.1%), while agricultural and pasture areas cover ∼ 25.8% (50). The study sites at La Huerta are located in the Chamela-Cuixmala Biosphere Reserve, a reserve created in 1993 that protects 1311.42 Km^2^ of well-preserved TDF and wetlands (51). The climate of the study area is predominantly warm subhumid with summer rains, with mean annual temperature ranging from 20 to 28 °C, and mean annual precipitation ranging from 600 to 2000 mm; the dry season goes from November to May (50). Several pollinator-dependent crops are cultivated during this season including watermelon (*Citrullus lanatus*), green tomato (*Physalis ixocarpa*), chili (*Capsicum annuum*), cucumber (*Cucumis sativus*), chayote (*Sechium edule*), papaya (*Carica papaya*), squash (*Cucurbita moschata* and *C. pepo*) and tomato (*Solanum lycopersicum*). These crops are usually located along river basins. Cultivation period is during the dry season as it allows farmers to reduce crop losses due to flooding and herbivory during the wet season (52,53). During 2017, the cultivated area for watermelon and green tomato in the three municipalities was 1,258 ha, and according to the SIAP, the total value production was 8,233,714 US dollars. The total income production of watermelon in the study area comprised on average 28.7% of the total income of agricultural production, and 7.3% of the cultivated area. Income production of green tomato comprised on average 6% of the total income of agricultural production and 7% of the cultivated area (54).

**Fig 1.**
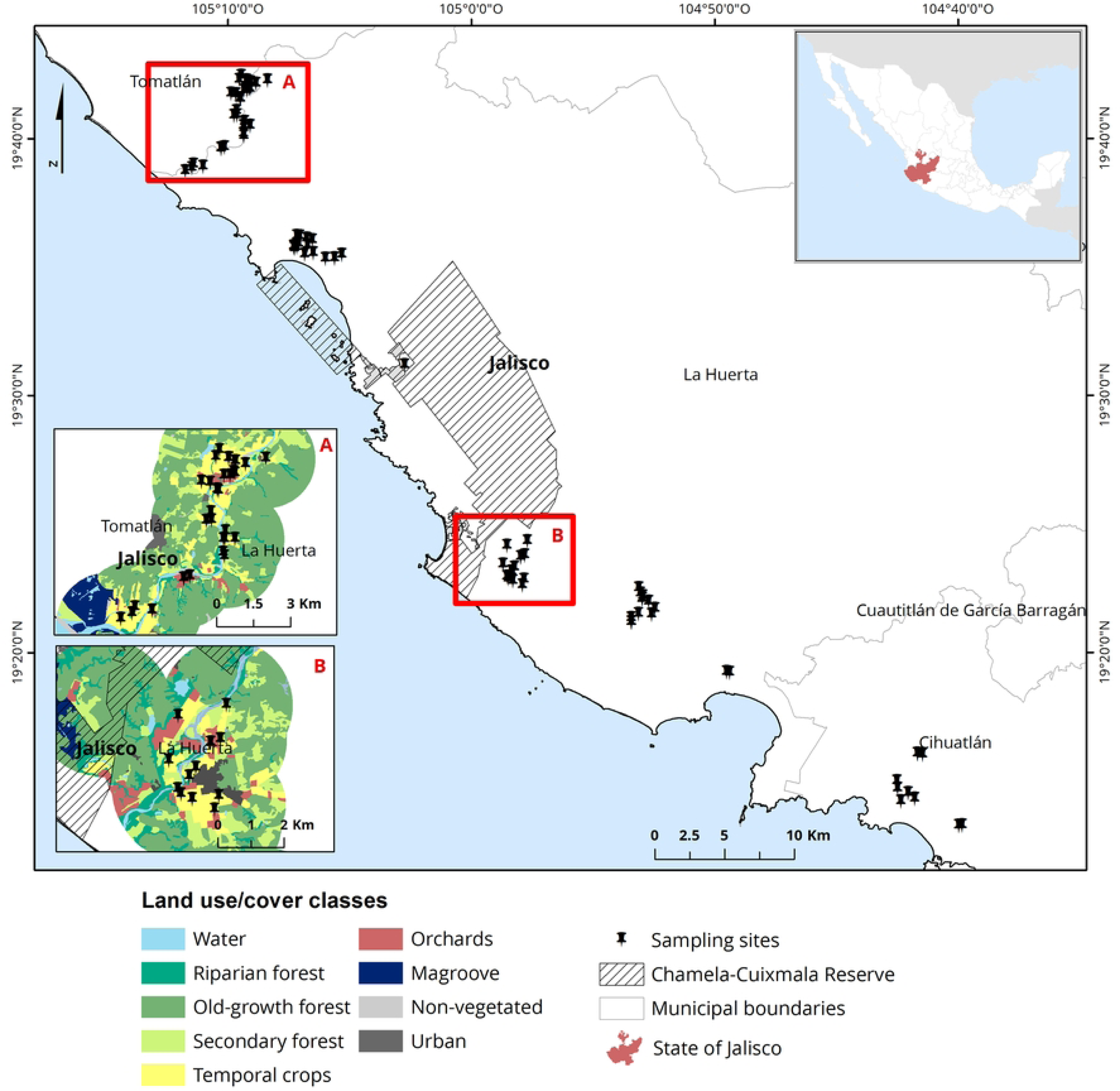
Study site. Study site and municipalities of La Huerta, Cihuatlan and Tomatlán where watermelon (*Citrullus lanatus*) and green tomato (*Physalis ixocarpa*) crops were sampled during the dry seasons of 2008, and 2014 to 2017. Letters A and B are examples of the land use/cover classes; A shows sampling sites distant from the Chamela-Cuixmala Reserve, and B shows sampling sites near the Chamela-Cuixmala Reserve.

### Study species crops

*Citrullus lanatus* (Watermelon): is native to Northeastern Africa and has been domesticated at least 4000 years ago (55). Flowers are solitary, 2-3 cm in diameter, with five light yellow petals. Watermelon is a self-compatible monoecious plant, with pistillate and staminate flowers (Fig 2), although some cultivars have hermaphrodite flowers (56); the species is pollinated by bees (1). In the study region, farmers use seedless watermelon as recipient plants and seeded watermelon as pollen donors in a 3:1 proportion, respectively. Seedles watermelon do not produce viable staminate flowers, they require to be pollinated with viable pollen from seeded plants (57–59). *Apis mellifera* is widely used as a pollinator of watermelon in the study region, where farmers have contracts with beekeepers to use *A. mellifera* hives to supplement watermelon pollination (at a cost of around $16 dollars per hive for the season during the study years).

**Fig 2.**
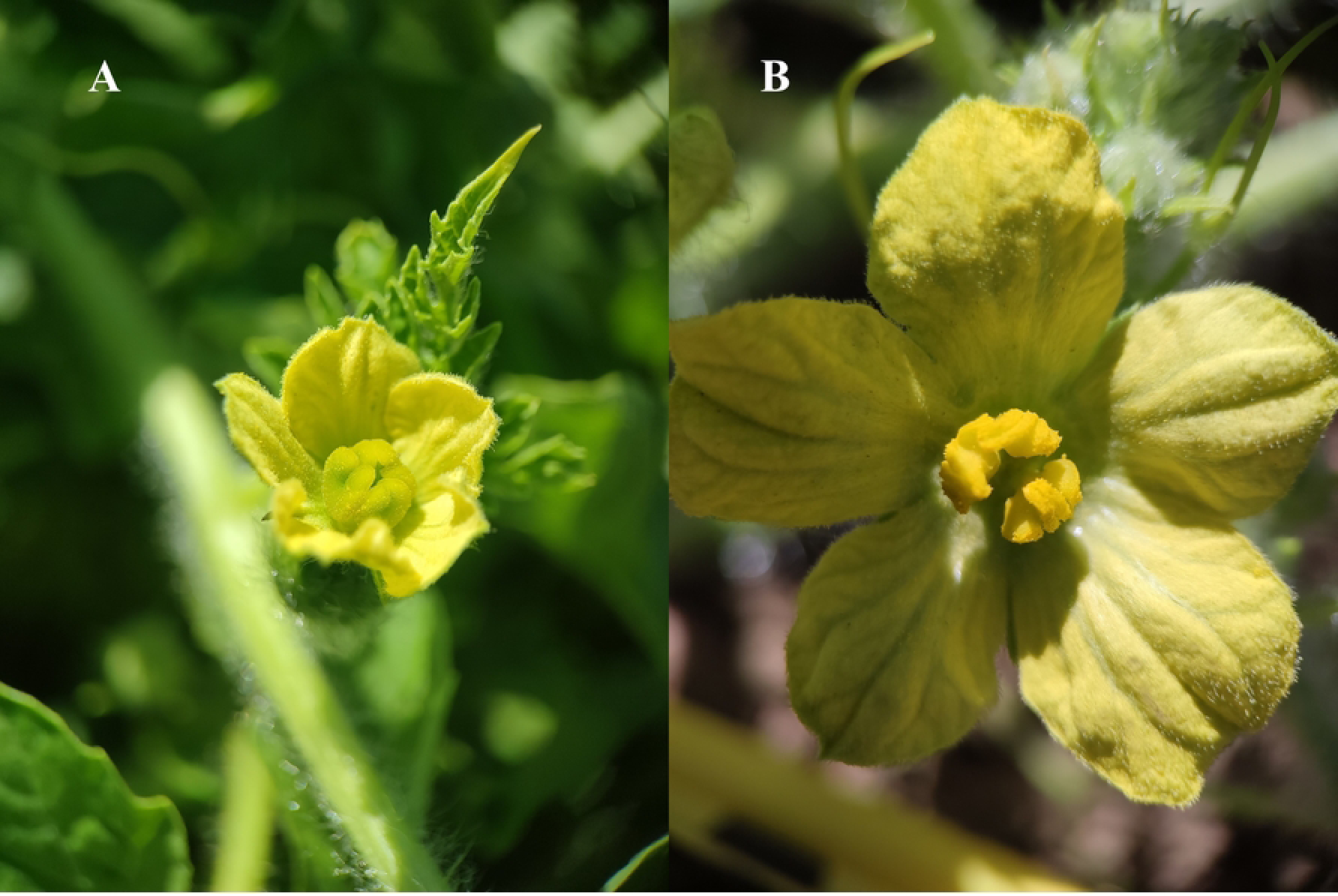
Flowers of Watermelon (*Citrullus lanatus*) from Jalisco, Mexico. (A) pistillate flowers and (B) staminate flowers.

*Physalis ixocarpa* (Green tomato): *Physalis* fruits have been used as a food source since the pre-Columbian era. There are about 90 species of *Physalis*, and of these, 70 species are found in Mexico (51–53). The main species of *Physalis* cultivated in Mexico is green tomato, the fifth economically most important vegetable. The flowers are hermaphrodite and solitary (Fig 3) and they open before anther dehiscence. The fruit is a fleshy green to yellow berry of variable size that is wrapped by a broad and persistent calyx. The species has been classified as self-incompatible in greenhouse studies (60–64).

**Fig 3.**
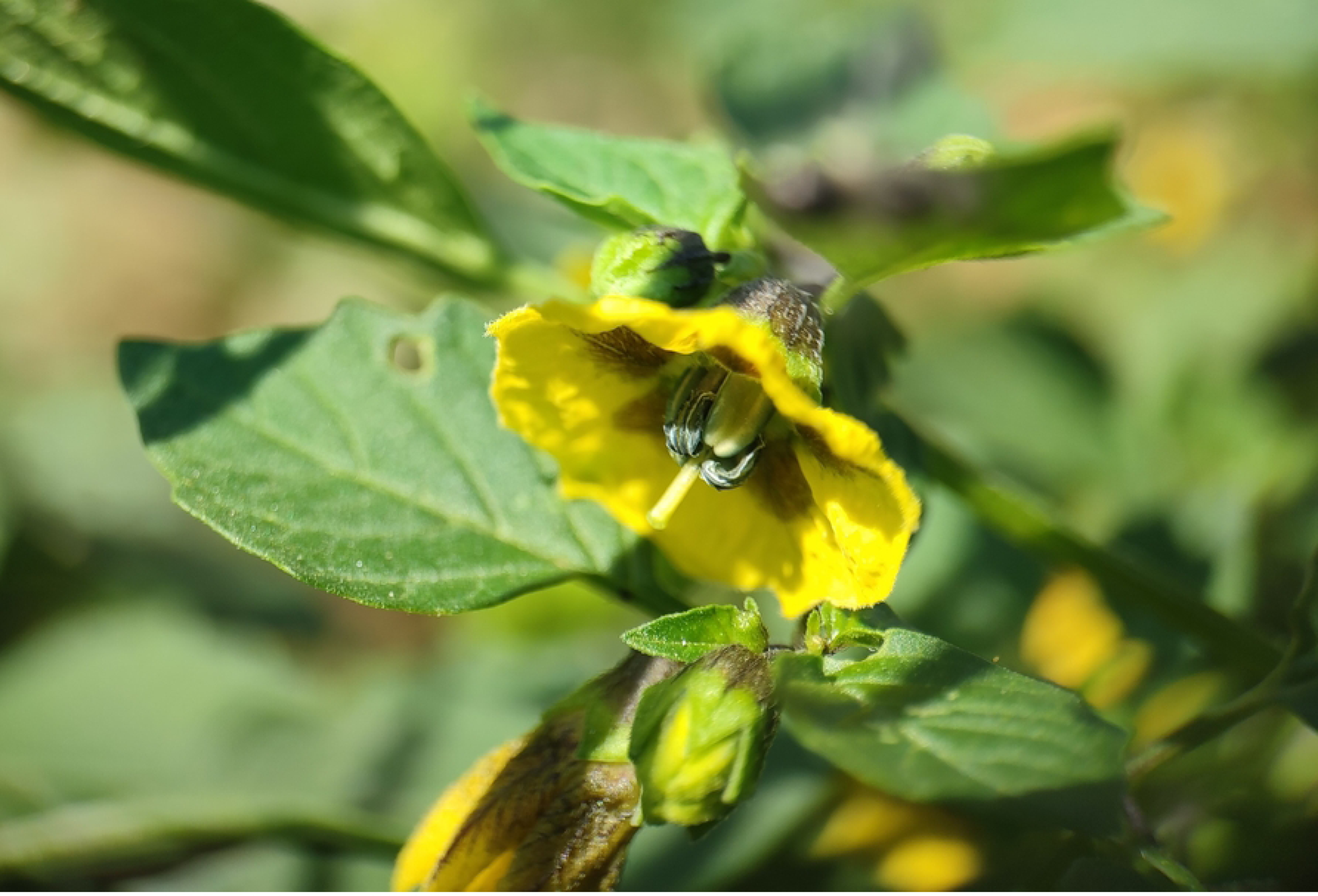
Hermaphrodite flowers of green tomato (*Physalis ixocarpa*) from Jalisco, Mexico.

### Study design

To evaluate pollinator effectiveness and the influence of the landscape on pollinator richness and abundance, over five years we sampled 39 plots of watermelon in 2008 and 2014-2017 (5, 11, 9, 6, 8 plots respectively), and 51 plots of green tomato in 2008, 2014-2017 (16, 14, 10, 7, 4 plots respectively). The number of crops varied in time because crop locations changed from year to year. To select the study plots for each crop, we searched the river basins of the study region in the early dry season each year. We selected plots where the study crops were at peak flowering (i.e., individual plants had a high proportion of flowers to fruits, about >0.8 of proportion). We georeferenced each surveyed plot. Pollinator community composition and visitation patterns, pollen load on pollinators bodies and pollinator dependence and pollinator efficiency experiments were recorded in years 2015-2017. Pollinator abundance data for the assessment of landscape and mapping crop pollination services using InVEST, were recorded in 2008 and 2014-2017.

### Pollinator community and visitation patterns

To characterize the community of floral visitors that are potential pollinators of the study crops, we surveyed flowers in seven watermelon plots for three years (2015, 2016, and 2017), and four green tomato plots for two years (2015 and 2016). We conducted video camera recordings in two to 10 plants per plot in one flower per plant from 900 to 1200 hrs.; each flower was recorded for 180 minutes. In watermelon plants, we surveyed both pistillate and staminate flowers. For both crops, we only registered floral visitors that contacted the reproductive organs of the flower. We recorded pollinator species or taxonomic group, time of arrival to the flower, and duration of the visit. We identified the pollinators visually when they arrived at the flower; when this was not possible, we captured the visitor with an insect net for further identification. We identified insects to the lowest possible taxonomic category with the help of taxonomic keys and field guides. We calculated visitation rates as the number of visits per flower per hour for each pollinator taxa. Time of arrival to the flower was used to obtain mean visitation rates by time of the day.

We conducted generalized linear mixed models with the GLIMMIX procedure in SAS version 9.4 (65) to evaluate differences among pollinator species (fixed effect) in pollinator visitation rates, and duration of individual pollinator visits (response variables). In the case of watermelon crops, we also evaluated how flower gender, pollinator species (fixed effects), and their interaction influenced the same response variables. We included year and field plot as random effects in both analyses. We specified a Poisson distribution and a log link function for both response variables. We specified the ILINK option of the LS-MEANS statement to obtain back-transformed least-square means and a Tukey adjustment for multiple comparisons.

### Pollen loads on pollinators’ bodies

To determine the capacity of different floral visitors to carry pollen of watermelon (*Citrullus lanatus*) and green tomato (*Physalis ixocarpa*), we captured insects visiting flowers (3-15 pollinator species) in the same study plots selected to document visitation frequencies in 2015. In the case of watermelon, we only captured pollinators in staminate flowers. We placed insects in separate vials and later removed pollen from each individual, dabbing one piece of fuchsin gel over four different parts of the body: back, head, ventral abdomen, and ventral torso. We did not remove pollen from specialized structures for pollen transport (i.e. corbiculae, scopae). We deposited each pollen sample on a slide and counted the number of pollen grains of watermelon using a stereoscopic microscope and the Zen program V 1.1.2 (66). Because of the difficulty to see green tomato pollen grains under a stereoscopic microscope, we counted grains in five regions of the slide with a magnification of 40X using an optical microscope and we averaged the number of pollen grains of each individual slide. To evaluate the capacity of different floral visitors to carry pollen of green tomato and watermelon, we performed a generalized linear mixed model with GLIMMIX procedure in SAS version 9.4 (65). The model included pollinator species as a fixed effect and the total pollen counts across the body or slide regions as a response variable. This analysis used a Poisson distribution and a log link function, the ILINK option of the LS-MEANS statement was used to obtain back transformed least square means, and field plot was included as a random effect in the model.

### Pollinator dependence and pollinator efficiency experiments

To determine the degree of dependence of watermelon (*Citrullus lanatus*) and green tomato (*Physallis ixocarpa*) on pollination by animals, in 2015 we conducted field experiments on 10 plants per plot on 11 plots for watermelon and 10 plots for green tomato. We marked two virgin flowers per plant that were assigned to one of two treatments: (1) open pollination, we marked flowers and left them exposed to visits by all potential pollinators; (2) pollinator exclusion, we covered flowers with meshnet bags one day before anthesis to exclude all floral visitors. On the third day after anthesis, we removed the bags and marked the flowers. For both treatments, we followed fruit development until maturation.

To determine the effectiveness of the most frequent floral visitors known to carry pollen of watermelon and green tomato, we added one more treatment in 2016 and 2017: (3) pollinator exclusion with one pollinator visit. We used 10 plants and one flower per plant and bagged the flowers one day before anthesis. We removed the bag the day after anthesis and allowed for a single pollinator visit. After the visit, we bagged the flower again and removed it three days after anthesis. We obtained data for individual visits of *Apis mellifera* and *Trigona fulviventris,* which were the most frequent pollinators.

Preliminary work suggested that green tomato produced fruits under pollinator exclusion; therefore, we also tested for apomixis adding an additional (4) emmasculation treatment to the green tomato plants evaluated in 2016 and 2017. One day before anthesis, one flower per plant was emasculated through the removal of all stamens using fine tweezers. The flowers were bagged until three days after anthesis and fruits were followed until maturity.

For both crop species, we compared fruit set among the pollination treatments. We performed generalized linear models with GENMOD procedure in SAS version 9.4 (65). The model used pollination treatment as the independent variable and the proportion of flowers that developed into fruit as a response variable. The analysis used a binomial distribution and a logit link function for fruit set. We used the Tukey-adjusted P-values for multiple comparisons.

We also tested the relationship between pollinator abundance (predictor variable) and fruit set (response variable) with a simple linear regression with the REG procedure in SAS version 9.4 (65). We used the insect abundance data from the transect surveys (see study design section) and the fruit set data from the open pollination treatments. The analysis used a binomial distribution and a logit link function for the variable fruit set.

### Influence of the landscape

To evaluate floral visitor abundance and richness at the landscape scale, we established 50 m random transects (one within each plot) and conducted flora visitor surveys in 2008 and 2014-2017. We sampled each plot only during sunny or partly cloudy days (when insects are more active), and at least after three days of any application of agrochemicals. We walked along each transect for 10 minutes registering all flower visitors that contacted the reproductive organs of the flowers. The insects were identified to the lowest possible taxonomic level in the laboratory with the help of taxonomic keys. To correlate the richness and abundance of pollinators of watermelon and green tomato to landscape configuration, we generated land-use and land-cover (LULC) maps for years 2007, 2012, and 2017, which provided the best quality maps to conduct analyses in close temporal proximity to the study years. LULC maps were elaborated through visual interpretation of SPOT5 satellite images at 5 and 10 m resolution from 2007, Rapideye satellite images at 5 m resolution from 2012, and Planet satellite images at 3 m resolution from 2017. Acquisition dates of satellite images were between February and May (the dry season when leaves fall off the trees in the tropical dry forest), corresponding closely to the sampling period of the bees. For the SPOT5 images, a fusion process was carried out between the 5-meter resolution panchromatic band and the multispectral bands to increase spatial resolution to 5 meters. To avoid spurious changes in LULC maps, we conducted an interdependent classification process, starting in 2007 and updated after 2010 and 2018. The satellite images were classified into nine LULC classes: (1) secondary forest (e.i., including pastures and forests in early successional stages), (2) deciduous tropical forest of advanced successional age (including intermediate and late successional stages), (3) riparian forest (including both gallery forest along large rivers and vegetation along the stream banks), (4) mangroves, (5) seasonal crops (e.g., corn, green tomato, tomato, chili, melon, watermelon, cucumbers, squash), (6) orchards (e.g., mango, papaya, coconut or citrus), (7) bare soil, (8) water, and (9) urban settlements. Image processing was done in ArcGIS 10.2 (67) and Qgis (68) at a scale of 1: 15 000. To validate LULC maps, we followed the methodology proposed by (69). We selected 553 random points from each LULC map, and then carried out verifications for each point using Google Earth images. For this procedure, we designated visual interpreters who had not been involved in the construction of the LULC maps. The overall accuracy of the classifications for 2007 and 2012 is 94%, while for 2017 is 96%. We obtained this calculation using the AccurAssess plugin for Qgis (70).

Around each polygon (i.e. plot) centroid, where the richness and abundance of pollinators were measured, we calculated circular buffer areas of 500m, 1000m, 1500, 2000m, and 2500m. Using the classified LULC maps for 2007, 2012, and 2017, we estimated the following parameters: (1) percentage of forest cover - including mangroves, tropical deciduous forest (secondary and late forest), and riparian vegetation-for each buffer area surrounding watermelon and green tomato crops, and (2) distance to the different forest types. Then, we assessed the correlation between the percent of forest cover and the distance to the different classes of forests *vs.* the following variables: (1) abundance and richness of *Apis mellifera*, (2) abundance and richness of native wild pollinator species, (3) total abundance. For each survey, we selected the LULC map closest to the field sampling year to estimate forest cover for each buffer area and to calculate Pearson’s correlation coefficient between forest areas and pollinator species abundance. Normality of residuals was assessed with the Shapiro-Wilk test. We conducted the spatial and correlation analysis using the free software environment R (71).

### Mapping crop pollination services using InVEST

To model crop pollination services provided by bee pollinators to Watermelon (*Citrullus lanatus* and *P. ixocarpa* crops, we used the crop pollination model implemented in InVEST (72). The model estimates an index of bee abundance across a landscape based on the availability of nesting sites (the different substrates and disponibility where bees can nest, in this study was cavity or ground substrates) and floral resources (plants with flowers that bee uses) within pollinator flight ranges as described below (72,73). The index of abundance produced by the model depicts the abundance of bees for each cell. The inputs required by the model are (a) a land use and land cover map, (b) land cover table describing LULC classes, and (c) a table describing guilds or species of pollinators to be modeled, their nesting biology and flight ranges (72,73). We used LULC map for 2017 (see “influence of the landscape” section) as the LULC input map. To estimate bee floral resources available in each LULC class, we conducted a literature review searching for plant species flowering from January to March in the TDF of the Chamela-Cuixmala Biosphere Reserve. We obtained the percentage of flowering plants from January to March found in each land cover class (S1). We used the list of pollinator species obtained from video recordings and through direct observation in sampled transects. We estimated each bee foraging distance from intertegular distance in mm using individuals captured for pollen counts. We recorded 2 - 20 measurements per species and calculated the typical foraging range (distance where 50% of foraging bees return to the nest, in social bees) in meters using the regression proposed by (74), this formula can applied to solitary bees (S2). We also reviewed the literature to determine nesting suitability (cavity or ground) for each bee species (S3). To compare INVEST predictions against the observed abundance of pollinators in watermelon and green tomato crops, we performed a simple linear regression using the REG procedure in SAS version 9.4 (65).

## Results

### Contribution of the main pollinators: Frequency and duration of visits

Watermelon (*Citrullus lanatus*): we observed eight species of bees visiting watermelon flowers. We found significant differences in pollinator visitation rates among species (F_(7,40)_ = 17, P ≥ 0.0001), and *A. mellifera* was the main visitor, accounting for 93.3% and 94.8% of total visits to pistillate and staminate flowers, respectively (Fig 4). Peak visitation occurred between 9:00 am and 11:00 am (Fig 5a). There were no significant differences in pollinator visitation rates by flower gender (F_(1,40)_ = 0.56, P = 0.45), nor for the interaction between flower gender and pollinator species (F_(5,40)_ = 1.27, P = 0.29). The average duration of a pollinator visit in flowers was 0.15±0.16 minutes. We did not find significant differences in the duration of a pollinator visit by flower gender (F_(1,1171)_ = 0.09, P = 0.77, Fig 6A), pollinator species (F_(5,1171)_ = 0.42, P = 0.8) or the interaction between flower gender and pollinator species (F_(4,1171)_ = 0.1, P = 0.98).

**Fig 4.**
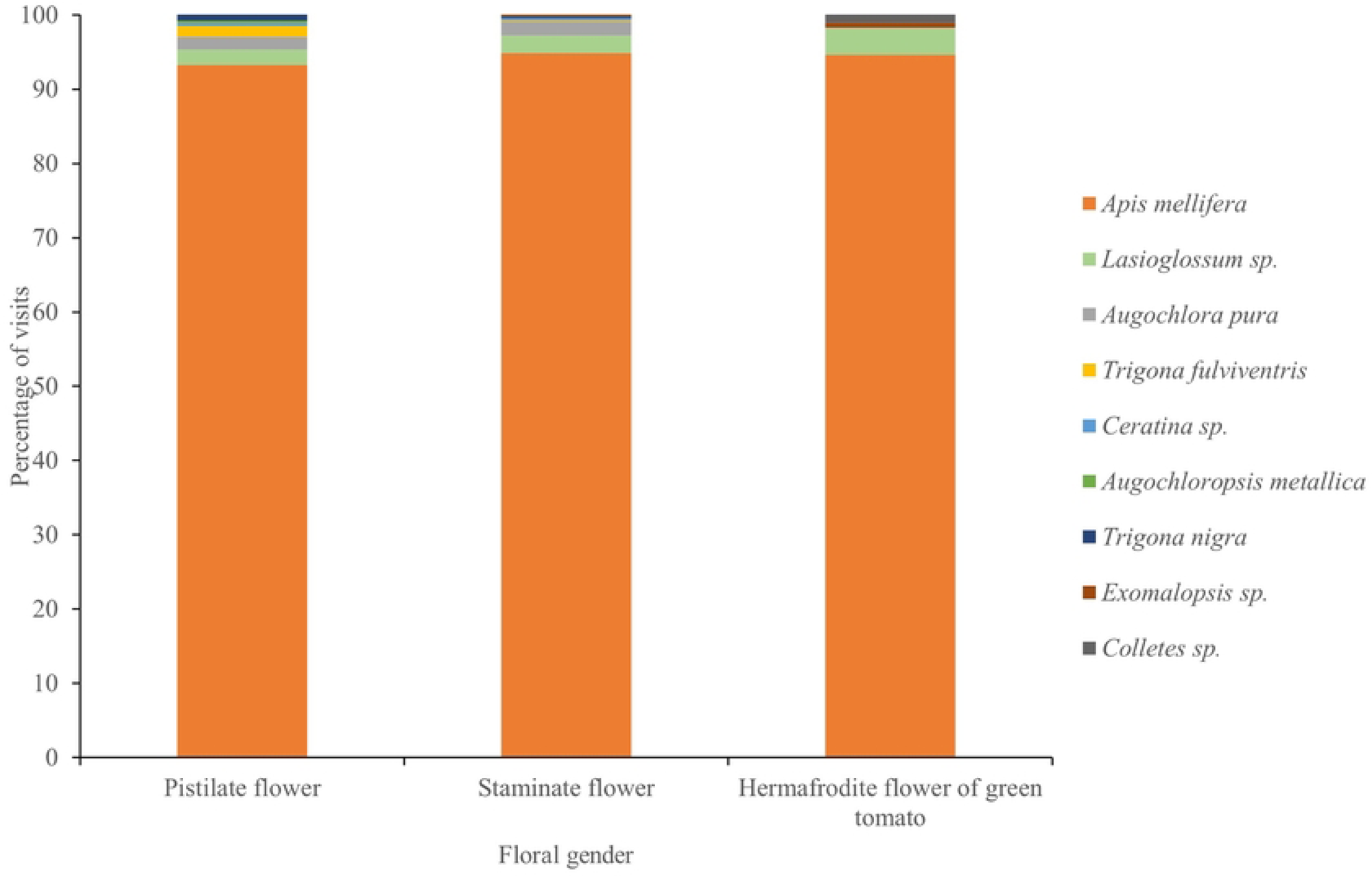
Composition and relative abundance of bee species visiting the male and female flowers of watermelon (*Citrullus lanatus*), and the bisexual flowers of green tomato (*Physalis ixocarpa*) in Chamela, Jalisco, Mexico across years 2015-2017.

**Fig 5.**
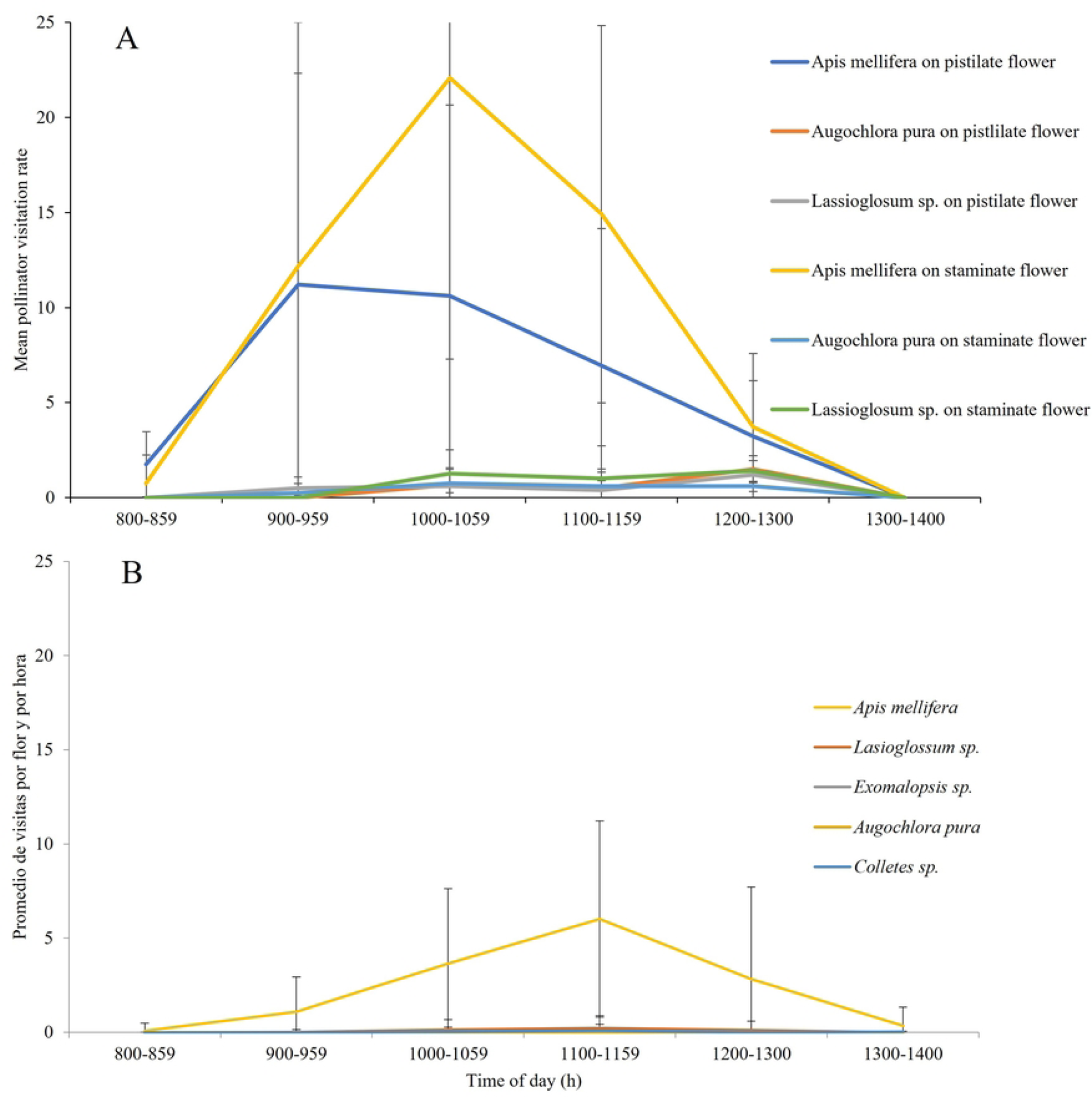
Pollination visitation rates (mean ± SE number of pollinator visits flower ^-1^ h ^-1^) to watermelon and green tomato flowers in Jalisco, Mexico during years 2015 to 2017. (A) Visits to watermelon male and female flowers, (B) visits to green tomato bisexual flowers. Different colors indicate different bee species and flower gender for the monoecious species watermelon (*Citrullus lanatus*).

**Figure 6.**
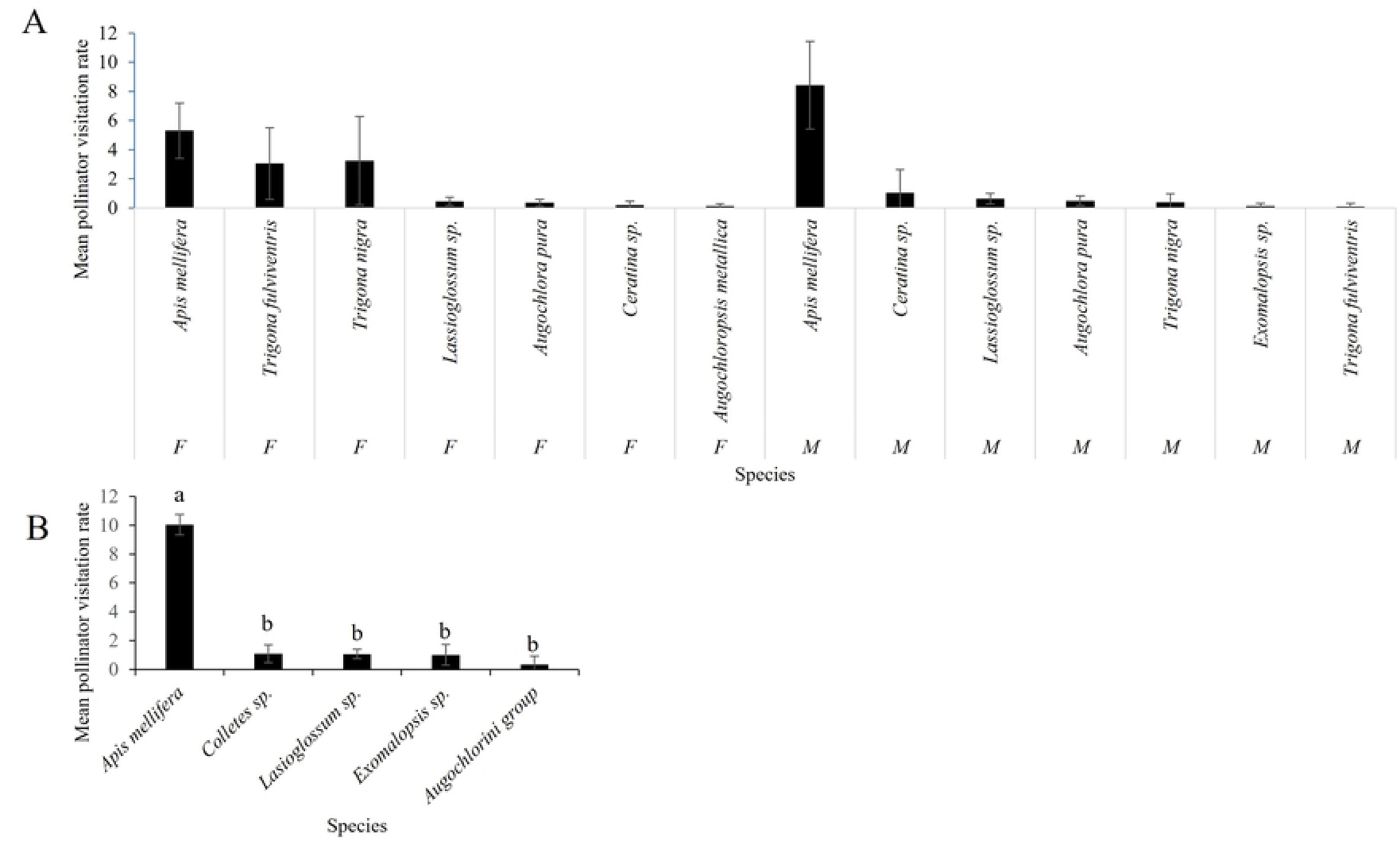
Pollinator visitation rates by bee species. Pollinator visitation rates (mean ± SE number of pollinator visits flower -1 h -1) for: (A) male (M) and female (F) flowers of watermelon (We did not find significant differences), and (B) bisexual flowers of green tomato (We find significant differences) in Jalisco, Mexico during years 2015 to 2017. For figure 6 B, different letters indicate significant differences between groups (*P* < 0.05) with Tukeýs ad hoc test.

Green tomato (*Physalis ixocarpa*): we observed five bee species visiting green tomato flowers. *Apis mellifera* was also the main visitor, accounting for 95% of total visits (Fig 4). Peak visitation rate occurred between 10:00 and 12:00 hrs (Fig 5b). We found significant differences in pollinator visitation rates by pollinator taxa (F_(7,33)_ = 156.42, P ≤ 0.0001; Fig. 6B). The average duration of pollinators in flowers was 0.15 ±0.12 minutes. We did not find significant differences in the duration of individual pollinator visits by pollinator taxa (F_(4,1008)_ = 0.22, P = 0.9).

### Contribution of the main pollinators: Pollen loads on pollinators’ bodies

Watermelon (*Citrullus lanatus*): We captured 40 individuals of seven bee species in watermelon male flowers. All captured bees had pollen on their bodies. We found significant differences in pollen count by pollinator taxa (F_(6, 21)_ = 592, P ≤ 0.0001; Fig. 7a). *Trigona fulviventris* and *Exomalopsis* sp. had the largest pollen grain loads, while halictid bees *Augochlora pura* and *Agapostemon* sp. had the smallest loads; *Trigona nigra*, *Apis mellifera* and *Lassioglosum* sp. carried intermediate pollen loads.

**Figure 7.**
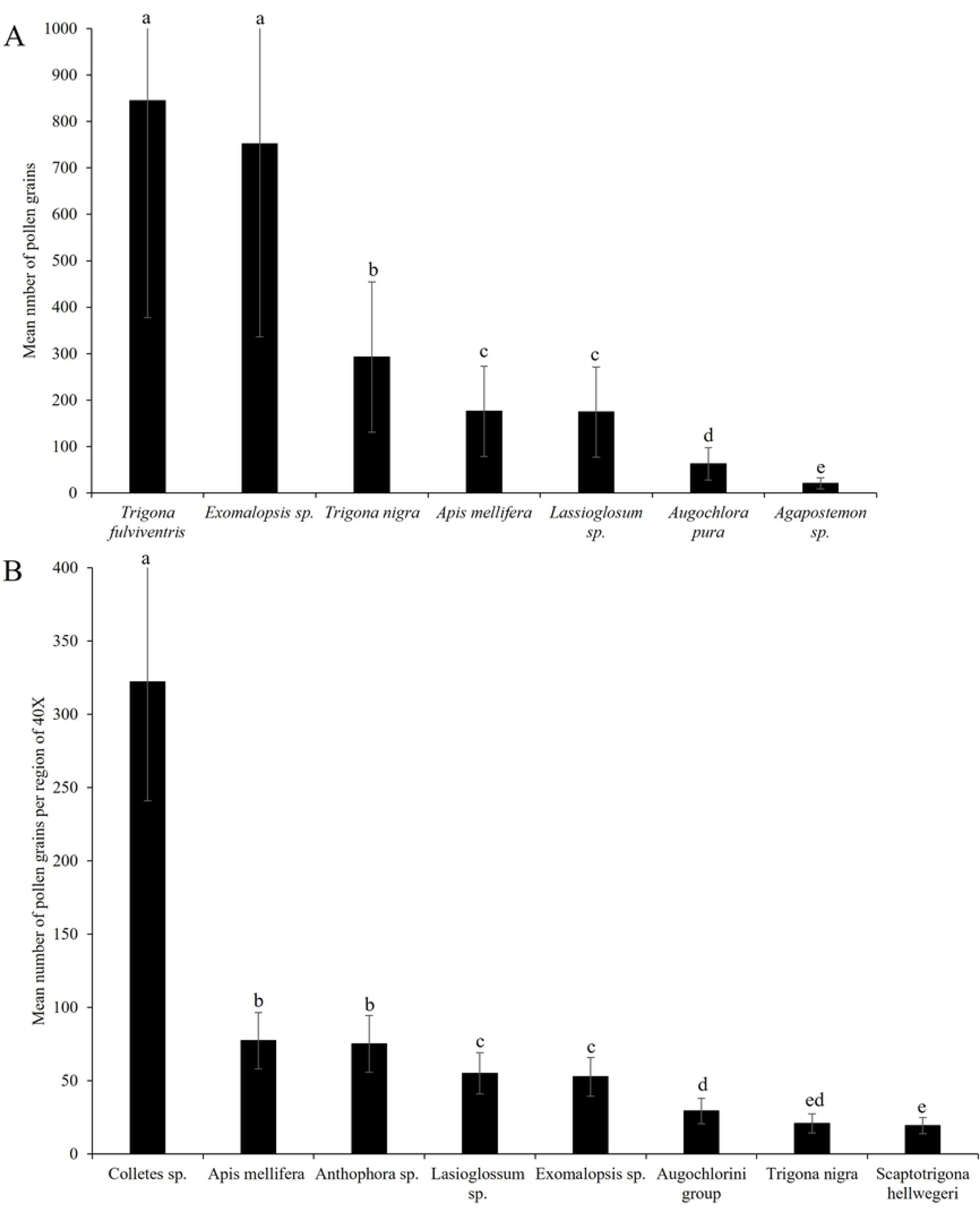
Pollen grains on the bodies of pollinators. Mean (± SE) number of pollen grains on the bodies of pollinator taxa collected in (A) Watermelon (*Citrullus lanatus*) and (B) Green Tomato (*Physalis ixocarpa*), in the region of Chamela, Jalisco, Mexico during years 2015 to 2017. Different letters indicate significant differences between groups (*P* < 0.05) with Tukeýs ad hoc test.

Green tomato (*Physalis ixocarpa*): We captured 45 individuals of eight bee species that had pollen on their bodies. We found significant differences in the number of pollen grains per load by pollinator taxa (F_(7, 33)_ = 156.42, P ≤ 0.0001; Fig. 7b).

*Colletes* sp had the largest pollen loads, *Apis mellifera*, *Anthophora* sp, *Lassioglosum* sp, and *Exomalopsis* sp carried intermediate loads, and bees of the Augochlorini group, *Trigona nigra* and *Scaptotrigona hellwegeri* carried the smallest pollen loads.

### Contribution of the main pollinators: Pollinator efficiency experiments on watermelon and green tomato

Watermelon (*Citrullus lanatus*): For the open pollination treatment 41 out of 142 marked flowers produced fruits, and fruit set was zero for the pollinator exclusion treatment. For the one pollinator visit treatment, out of 20 flowers with one visit of *Apis mellifera*, no fruits were produced. The regression between pollinator abundance and open fruit set was not significant, less than one percent of the variation in fruit set was explained by pollinator abundance (F_(1, 9)_ = 0.07, P = 0.8, *R^2^* = 0.007, Fig. 8A).

**Figure 8.**
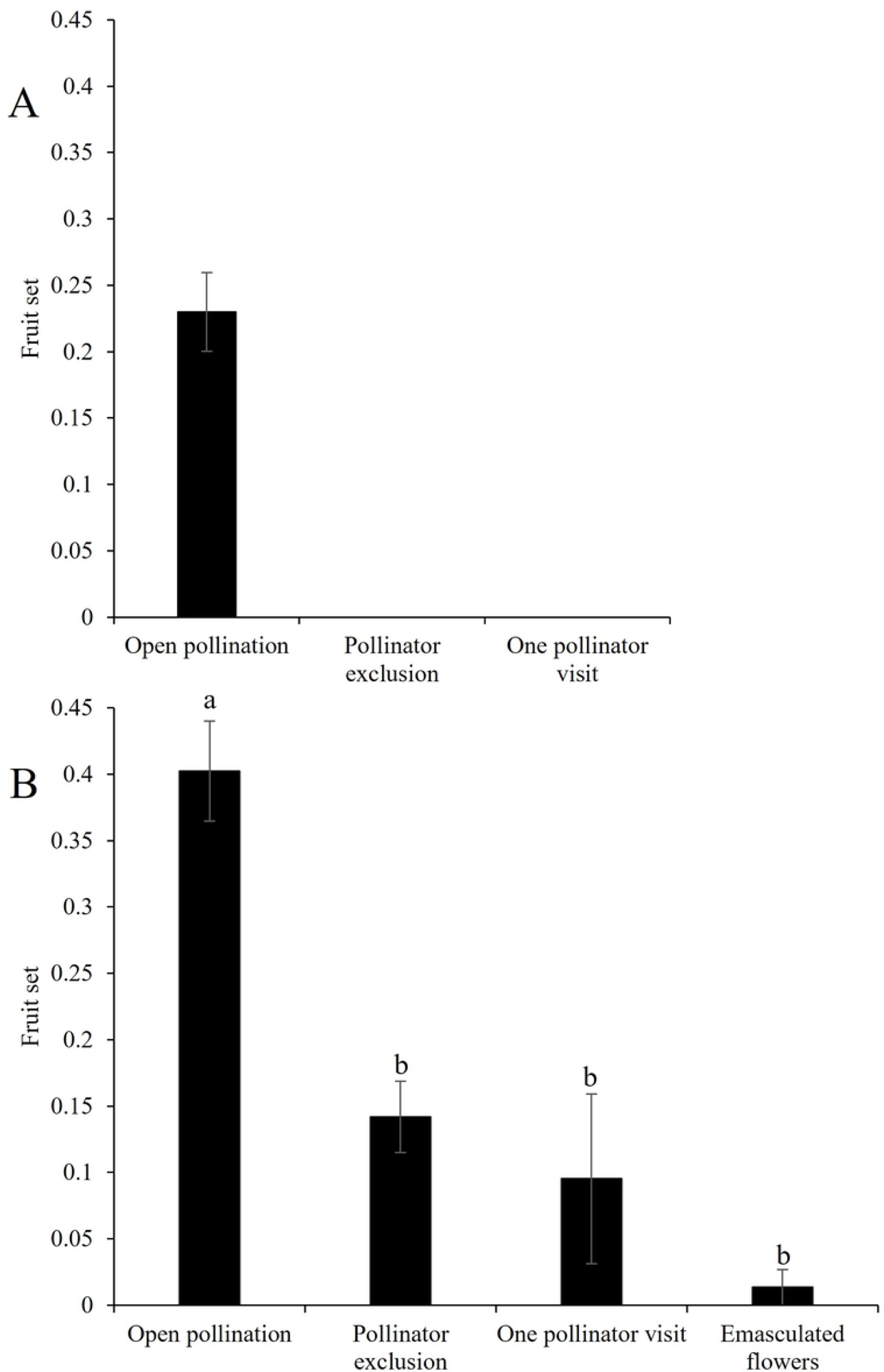
Pollination efficiency treatments. Mean (± SE) fruit set of pollination efficiency treatments conducted to determine the potential for autonomous fruit production (in the absence of pollinators) and the pollination efficiency of *Apis mellifera* (one pollinator visit). (A) Watermelon (*Citrullus lanatus*) (B) Green tomato (*Physalis ixocarpa*) plants. Different letters indicate significant differences between treatments (p < 0.05) with Tukey ad hoc test.

Green tomato (*Physalis ixocarpa*): For the open pollination treatment, 68 out of 169 flowers produced fruit. For the pollinator exclusion treatment, we obtained 19 fruits out of 167 flowers. For the one pollinator visit treatment, we only obtained two fruits from 21 flowers. For the emasculation treatment conducted to assess for apomixis, we only obtained one fruit from 74 flowers. We found significant differences between treatments, where open pollination was higher than the other treatments, and in the absence of pollinators, green tomato crops reduced their fruit production by 65% (ꭓ^2^ = 65.94, df = 3, P ≤ 0.0001; Fig 8B). We found a significant regression between pollinator abundance and the fruitset of open-pollinated flowers (*R*^2^= 0.68; F_(1, 8)_ = 17.63, P = 0.003).

### Influence of the landscape

For watermelon (*Citrullus lanatus*) crops, in five years we observed 15 pollinator species in 39 plots (S4). For green tomato (*Physalis ixocarpa*) crops, in five years we observed 19 species of pollinators in 51 plots (S4). In crops of green tomato for year 2008, we found significant positive regressions where forest cover predicted the total abundance of pollinators at the following distances to the focal crop: 500m (*R*^2^=0.5, P = 0.003), 1000m (*R*^2^ =0.5, P = 0.001), 1500m (*R*^2^ =0.55, P = 0.001), and 2500m (*R*^2^ of 0.5, P = 0.003) (Fig. 9). We did not find significant differences with distance to the forest and forest cover in 2014 or 2016. For watermelon, we did not find significant effects of landscape configuration on pollinator diversity.

**Figure 9.**
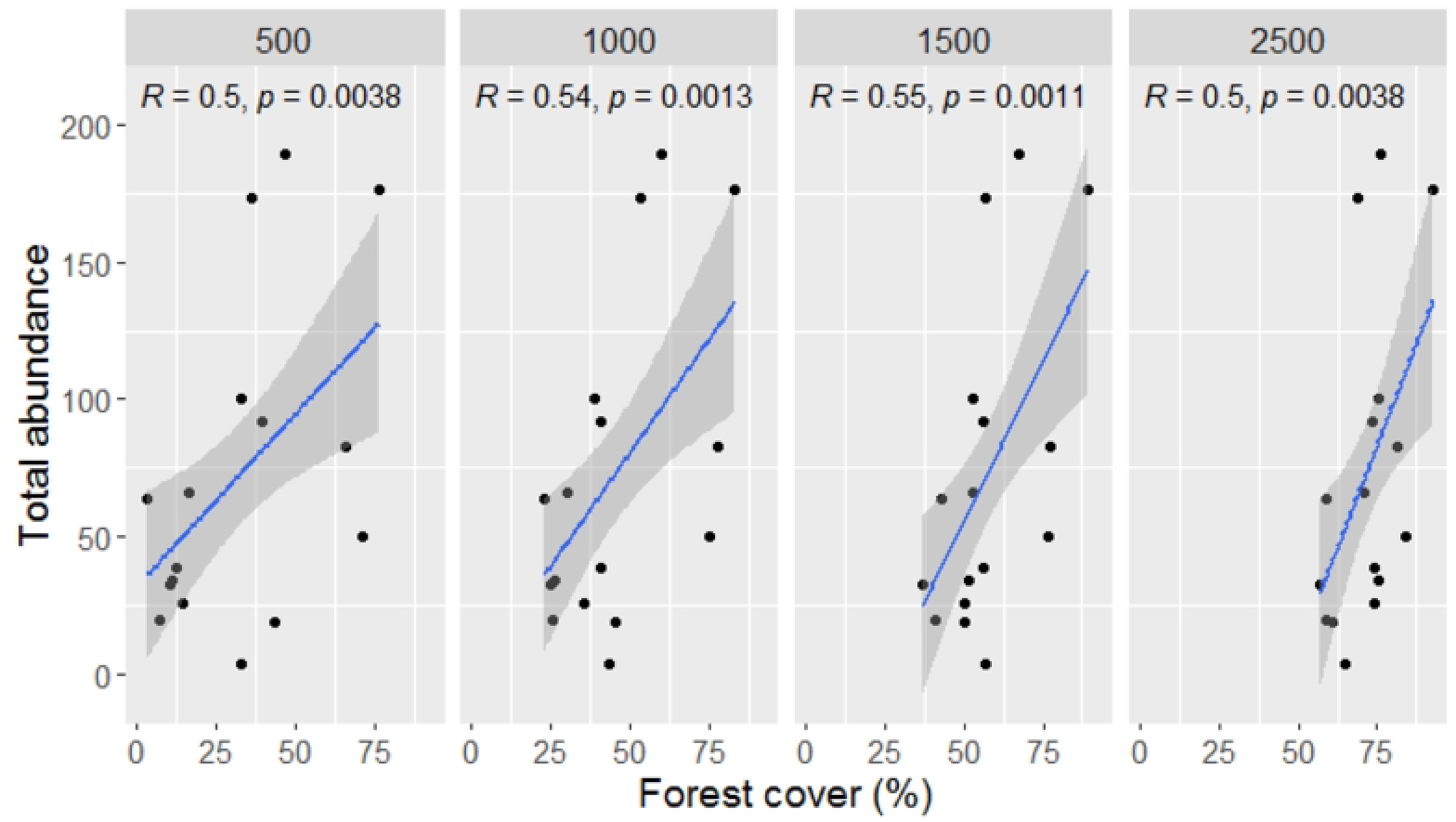
Correlation between the abundance of pollinators in Green tomato (*Physalis ixocarpa*) crops and forest cover in 16 crops for year 2008.

### Use of InVEST Natural Capital Project Program

We obtained the per-pixel total abundance of 12 pollinators species for three years. We found a significant positive regression between the pollinator abundance predicted by the model for 2008 and: (1) the observed total pollinator abundance (F_(1, 19)_ = 5.41, *R*^2^=0.22, P = 0.03), (2) the observed abundance of social bees (F_(1, 8)_ = 5.37, *R*^2^ =0.22, P = 0.03). We did not find a strong regression between the observed abundance and the predicted abundance of INVEST for 2014 and 2016 pollinator abundance.

## Discussion

### Pollination of watermelon and green tomato crops in western Mexico

Watermelon (*Citrullus lanatus*) and green tomato (*Physalis ixocarpa*) are two economically important crops for Mexico, the first one native to Africa and the second one native to Mesoamerica. The results of this study showed that, despite their different geographic origins and floral morphologies, the honeybee *A. mellifera* is the main floral visitor, accounting for more than 90% of the visits to flowers of both species. Honeybees visited flowers throughout floral anthesis and in terms of pollen carrying capacity, they were preceded only by a native bee species in the genus *Colletes*, a rare floral visitor in both crops. Honeybees are key pollinators of many crops worldwide (8,21,75), and part of their success is related to their social behavior, their high reproductive rates, and their ability to establish large populations and feed on diverse floral resources (21,76,77). Moreover, africanized honeybees have successfully established populations across diverse habitats in the New World, including natural ecosystems, agricultural land and urban areas (21,22,78). In the Chamela-Cuixmala Reserve, africanized honeybees have established feral populations interacting with native plants and neighboring crops (19,22). The high visitation frequency and pollination effectivenesss of *A. mellifera* make this bee species the most important provider of pollination services for watermelon and green tomato in the study region.

A key trait of honeybees as pollinators of watermelon and green tomato is their ability to forage throughout the year, even in seasonal environments, such as the Chamela-Cuixmala region (19,22). Moreover, africanized honeybees can locate and exploit floral resources in times of limited floral availability, and they may travel large distances to find food (77,79,80). These behaviors enable them to sustain colonies during the dry season. In contrast, native bees include both social and primitively solitary species that have smaller populations and foraging ranges (80), particularly during the dry season. Eight species of native bees were observed visiting the flowers of both crops at low frequencies in flowers, but some of them were able to carry high pollen loads on their bodies. These included social *Trigona fulviventris* and solitary *Exomalopsis* sp for watermelon, and solitary *Colletes* and *Anthophora* sp for green tomato crops (Fig. 4). Native bees can be effective pollinators of watermelon crops in other regions, as was demonstrated for the sweat bee *Lassioglosum* sp. in Pakistan, and for *Bombus impatiens* in USA (75,81). However, in our study system, native bees were found in low abundance, possibly due to the time of cultivation of watermelon and green tomato. It has been demonstrated that the abundance and diversity of flowers and floral visitors experience a notable reduction in the dry season in the Chamela-Cuixmala region (22). During the rainy season in this region or in less seasonal ecosystems, native bees and other pollinators may play an important role in the pollination of commercial crops, even in places where honeybees are dominant (8,21,82,83).

The results of this study suggest that, in order to reach the natural fruit set observed in watermelon (23%) and green tomato (40%), both plant species require multiple pollinator visits. In this context, native bees likely play a complementary role in the pollination of these crops. Research has shown that there is often complementarity between *A. mellifera* and native pollinators, which can enhance crop pollination; this complementary effect has been observed in various crops, including almonds and sunflowers (24,25). A previous study found that honey bees were the most abundant and important pollinators of squash crops (*C. moschata*) during the dry season, when native pollinators are not present; however, during the rainy season, native bees of the genus *Peponapis* were the most important pollinators of squash crops due to their abundance and greater effectiveness (19). Similarly, the native pollinators of wild *Physalis* species are only active during the rainy season and they are oligolectic, meaning that they feed on a small range of plant species. For example, the main pollinator of *Physalis viscosa* is the social stingless bee *Perdita halictoides*, which feeds primarily from species of the genus *Physalis* (84). Therefore, the study of wild crop relatives and their interaction with native bees is crucial to the understanding of crop pollination dynamics.

In addition to assessing the interactions between crops and their native and managed pollinators, it is essential to evaluate the extent to which a crop relies on pollinators for reproduction. Our results showed a 65% decrease in green tomato yield in the absence of pollinators when contrasted with open pollination (Fig. 8), suggesting the species is highly dependent on pollinators for reproduction.

However, despite being a self-incompatible species, fruit set is possible in the absence of pollinators in *P. ixocarpa,* suggesting a possible breakdown of the self-incompatibility reproductive system towards the last days of anthesis in the absence of cross-pollination. Our study revealed that green tomato flowers have a lifespan of three days; however, the anthers become dehiscent on the second and third day, enabling delayed self-pollen deposition. This process has been described for other Solanaceae species with plastic self-incompatibility systems. In *Nicotiana alata* and *Solanum carolinense*, the decay of the RNAses involved in the self-incompatibility system occurs before the end of anthesis if the flowers are not previously cross-pollinated. This phenomenon suggests the potential for reproductive assurance through self-pollination (85,86).

### Assessment of landscape influence on pollinator diversity and abundance

The conservation condition and distance to natural forests are determinant factors that ensure the richness and abundance of insects; therefore, the proximity of crops to continuous forests is expected to increase pollination service to agricultural systems (29,41). Moreover, landscapes that incorporate large expanses of natural habitat exhibit a greater diversity and abundance of pollinating insects and a corresponding increase in pollination services compared to landscapes with scant natural habitat (86). In the case green tomato (*Physalis ixocarpa*) crops, which are pollinated by honeybees present in the wild-agricultural interface, we observed a positive relationship between pollinator abundance and forest cover for year 2008. Other studies have demonstrated that proximity to continuous natural habitats enhances pollination services to agricultural fields by increasing pollinator abundance, visitation and diversity, pollen deposition, fruit set, and crop productivity (30,39–41,87,88). However, the relationship between forest cover and pollinator abundance and diversity was not significant for green tomato during 2014-2017. This could be related to the recent rise in storm intensity associated with global climate change, which in the region of Chamela, was evidenced by the direct impact of two large hurricanes, Jova in 2011 (category 3) and Patricia in 2015 (category 5). The impacts of hurricanes was evidenced by dramatic changes in vegetation greenness (89) and across extensive areas of fallen trees and branches across the Chamela-Cuixmala Biosphere Reserve (89,90). These events possibly created a reduction in the availability of nesting cavities and floral resources for bees.

In the case of watermelon, no significant association between insect abundance and distance from the forest was found in any year for watermelon. As in green tomato, the passage of hurricanes reducing insect populations and changing landscape structure may partly explain this result. However, the lack of an association between landscape traits and pollinator abundances, may also be explained by the fact that managed honeybee hives were used to supplement pollination and increase crop yield in watermelon fields during the study years. As previously mentioned, highly social bees like *A. mellifera*, have large populations and extensive foraging ranges; this enables them to exploit floral resources from agricultural fields located at significant distances from natural habitats (77,79).

## Conclusions

In the central pacific coastal region of Jalisco, Mexico, pollinator dependent crops like watermelon and green tomato are important for regional and local economies. Natural ecosystems can be a reservoir of pollinators, including both native species and feral honey bee populations; thus, changes or lack of natural ecosystems and thus pollinators can impact agricultural production and the economy of farmers in the region. The maintenance of natural areas and riparian corridors will be key to the conservation of pollinators and pollination services in natural and agroforestry ecosystems.

## Acknowledgement

We thank two anonymous reviewers for providing feedback on the manuscript. We also thank E. Perez, E. Paramo, G. Garcia, J. Cristobal, A. Braga, H. De Santiago, F. Parraguirre for laboratory and field assistance, and Gumersindo Sanchez-Montoya for technical assistance. We thank the farmers of the region Costa Alegre who supported this study.

